# High-throughput and scalable single cell proteomics identifies over 5000 proteins per cell

**DOI:** 10.1101/2023.11.27.568953

**Authors:** Zilu Ye, Pierre Sabatier, Leander van der Hoeven, Teeradon Phlairaharn, David Hartlmayr, Fabiana Izaguirre, Anjali Seth, Hiren J. Joshi, Dorte B. Bekker-Jensen, Nicolai Bache, Jesper V. Olsen

## Abstract

The emergence of mass spectrometry (MS)-based single-cell proteomics (SCP) promise to revolutionize the study of cellular biology and biomedicine by providing an unparalleled view of the proteome in individual cells. Despite its groundbreaking potential, SCP is nascent and faces challenges including limited sequence depth, throughput, and reproducibility, which have constrained its broader utility. This study introduces key methodological advances, which considerably improve the sensitivity, coverage and dependability of protein identification from single cells. We developed an almost lossless SCP workflow encompassing sample preparation to MS analysis, doubling the number of identified proteins from roughly 2000 to over 5000 in individual HeLa cells. A comprehensive evaluation of analytical software tools, alongside strict false discovery rate (FDR) controls solidified the reliability of our results. These enhancements also facilitated the direct detection of post-translational modifications (PTMs) in single cells, negating the need for enrichment and thereby simplifying the analytical process. Although throughput in MS remains a challenge, our study demonstrates the feasibility of processing up to 80 label-free SCP samples per day. Moreover, an optimized tissue dissociation buffer enabled effective single cell disaggregation of drug-treated cancer cell spheroids, refining the overall proteomic analysis. Our workflow sets a new benchmark in SCP for sensitivity and throughput, with broad applications ranging from the study of cellular development to disease progression and the identification of cell type-specific markers and therapeutic targets.

## Main

Multicellular organisms are comprised of specialized tissues that consist of various cell types, each performing distinct functions. The unique properties of each cell type arise from the interaction of their genetic information with internal factors. Studying individual cells is critical since cells can exhibit diverse behaviors under identical external environments^1^. Single-cell RNA sequencing has transformed cell biology by providing a granular view of gene expression patterns, spatial cell architecture, cellular heterogeneity, and dynamic cellular responses^2^. However, mRNAs serve as an intermediate step in gene expression and does not directly reflect cellular activity. Studying proteins provides a more direct and comprehensive understanding of cellular functions, regulatory mechanisms, and disease processes compared to studying mRNA changes alone. Protein analysis captures post-translational modifications, protein diversity, and functional aspects that are not reflected in mRNA analyses.

Single-cell proteomics (SCP) is emerging as the next frontier in proteomics and has already enhanced our understanding of cellular differentiation and diseases by allowing for the direct measurement of single-cell proteomes and their post-translational modifications (PTMs)^3-7^. This capability is instrumental in delineating the functional phenotypes within cell populations, elucidating cellular and embryonic development, forecasting disease trajectories, and pinpointing specific surface markers and potential therapeutic targets unique to each cell type^8, 9^.

SCP predominantly utilizes two main quantitative techniques: label-free^5, 10, 11^, or multiplexed^6, 7, 12^ analysis. Label-free quantitative (LFQ) analysis presents the simplest SCP workflow by lysing cells to extract proteins, which are subsequently digested into peptides, separated, and analyzed using mass spectrometry (MS), thus identifying and quantifying peptides from single cells. Multiplexed analysis, on the other hand, employs isobaric labeling, allowing for the simultaneous analysis of multiple samples by chemically tagging peptides with unique stable isotope encoded mass tags. The recent innovation of non-isobaric multiplexed Data Independent Acquisition (plexDIA^13^ or mDIA^14^) has combined the strengths of DIA with multiplexing, improving protein quantification rates and accuracy without the ratio compression problems associated with Tandem Mass Tags (TMT)^15-17^. State-of-the-art single-cell proteomics, identifying around 1000–2000 protein groups per cell and 1500–2500 proteins across cells, required improvements in MS sensitivity and sample preparation^10^. Yet, the loss of peptides during sample preparation and analysis due to protein adsorption loss, chemical modifications, and ion manipulation remains a challenge^18^.

To address these challenges, we developed a nearly lossless LFQ-based SCP method, which identifies over 5000 proteins and 40,000 peptides in single HeLa cells. Our workflow involves single cell dispensing and sample preparation using the cellenONE with a proteoCHIP EVO96 and direct transfer to Evotip disposal trap columns, and subsequent analysis using the Evosep One LC with Whisper flow gradients coupled to narrow-window data-independent acquisition (nDIA) on the Orbitrap Astral mass spectrometer^19^ (**Fig. 1a, Supplementary Video 1**). Our study includes a systematic evaluation of database search tools and an error-rate estimation using an entrapment approach, ensuring reliable data analysis. Furthermore, our method has enabled the direct investigation of PTMs in single-cell proteomes, achieving deep coverage in phosphorylation and glycosylation without prior enrichment. The application of this technique to spheroid samples with a new dissociation buffer underscores its robustness, offering significant insights into the proteomic intricacies of individual cells and their implications for biological processes and disease states.

**Figure 1.**
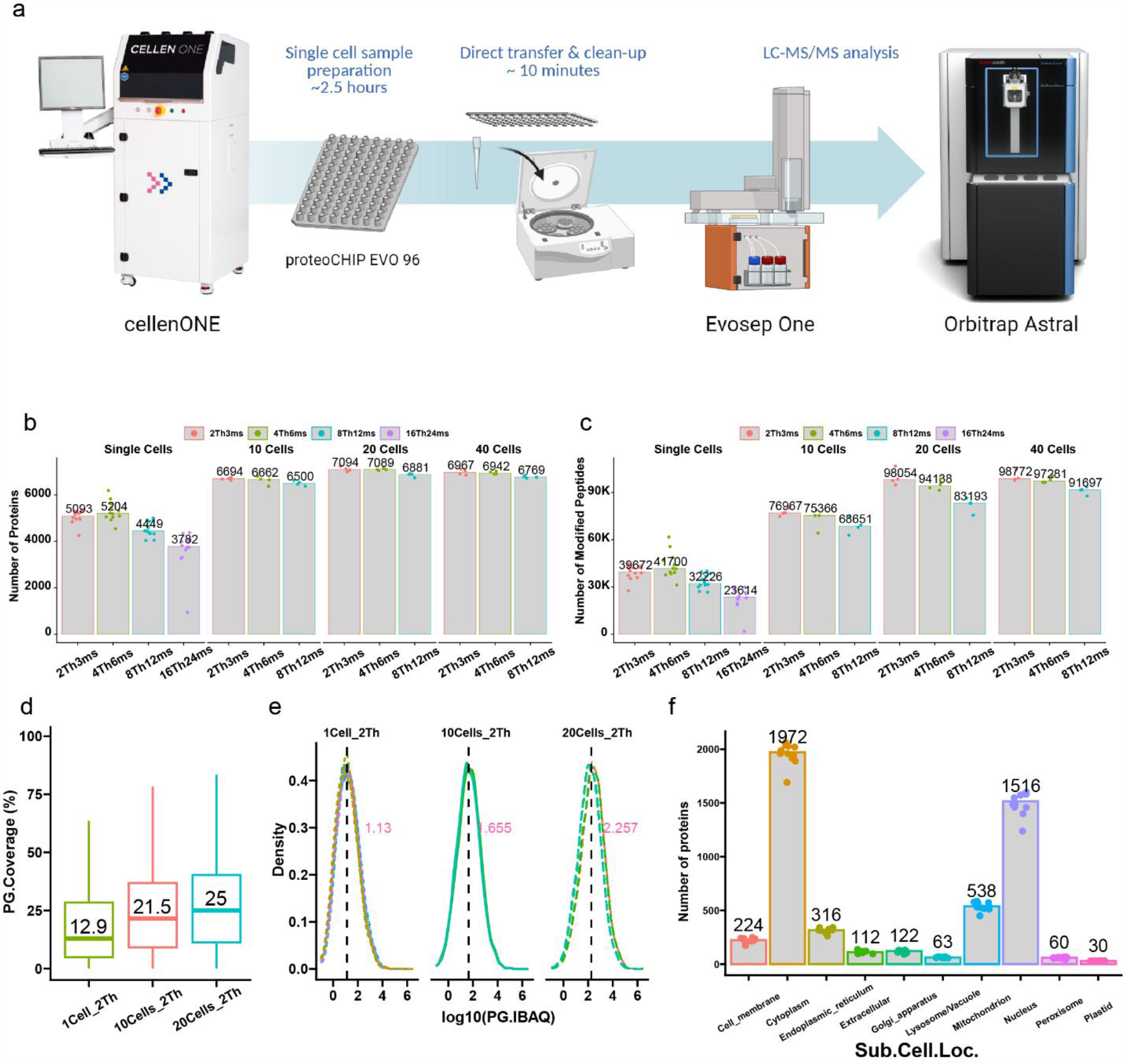
Workflow and Results of High-Throughput Label-Free Single-Cell Proteomics. (**a**) Schematic of the single-cell isolation and proteomic processing workflow. (**b**) Number of proteins identified in single, 10, 20 and 40 HeLa cells using variable nDIA settings. (**c**) Number of peptides identified in single, 10, 20 and 40 HeLa cells using variable nDIA settings. (**d**) Protein sequence coverage percentages for single-cell, 10-cell and 20-cell samples. (**e**) Distribution of iBAQ values showing quantification range and consistency across different cell counts. (**f**) Overview of protein localization across cellular compartments. Spectronaut (v18) was used for the search.

## Results

### Roadmap to high-throughput label-free proteomics with great depth

Key considerations for single-cell proteomics sample preparation workflows include minimizing surface adsorption losses by keeping samples as concentrated as possible, and reducing buffer evaporation and pipetting steps for reproducibility and maximizing detection sensitivity. Consequently, we focused on developing a SCP workflow characterized by ultra-high sensitivity (**Fig. 1a, Supplementary Video 1**). Initially, cells were isolated and processed using a one-pot technique in the cellenONE X1 platform (**Supplementary Figure 1a, 1b**). A key innovation is the proteoCHIP EVO 96, designed for single-cell sample preparation, which operates with minimized volumes at the nanoliter level, enabling the simultaneous proteomics sample preparation of up to 96 samples in parallel (**Supplementary Figure 1c**). This chip is precisely tailored for compatibility with the Evosep One LC system for a streamlined sample transfer process that is free from additional pipetting steps. For analysis by liquid chromatography tandem mass spectrometry (LC-MS/MS), we combined the Whisper flow methods on Evosep One with high-precision IonOpticks nanoUHPLC columns to maximize sensitivity (**Supplementary Figure 1d**). Our recently introduced narrow-window DIA (nDIA) method^19^, applied on the Orbitrap Astral mass spectrometer, significantly amplifies the sensitivity and efficiency of the SCP analysis. To optimized the nDIA approach for the Orbitrap Astral^20, 21^, which had not been previously used for label-free single-cell analysis, we initially assessed different nDIA methods, examining different quadrupole isolation windows and scaled ion injection times (IT) for DIA-MS/MS scans, accordingly. Through this comparative analysis, we determined the optimal nDIA method and found that using 4-Th DIA windows and 6-ms max IT (4Th6ms) resulted in the highest proteome coverage, leading to the identification of a median number of 5204 proteins in single HeLa cells and >6000 proteins in one of the HeLa cell preparations (**Fig. 1b**). We observed that as the IT increased in the 8Th12ms and 16Th24ms methods, there was a corresponding decline in the number of proteins identified, likely attributable to an increase in chemical noise signals within the analyzer. When we expanded our method to process larger cell quantities, we identified over 7000 proteins from a batch of only 20 cells. The achieved depth at the peptide level was particularly remarkable, with median identifications of 41,700 peptides for single-cell samples and 98,054 peptides for samples from 20 cells (**Fig. 1c**), resulting in median protein sequence coverage of 12.9% for single cells and 25% for 20-cell samples (**Fig. 1d**). This profound peptide coverage facilitated highly accurate protein quantification, with intensity Based Absolute Quantitation (iBAQ) values exhibiting an extensive dynamic range that spanned ∼ five orders of magnitude (**Fig. 1e, Supplementary Figure 2**). The relative protein abundance estimates also demonstrated an almost linear relationship of iBAQ values across samples with varying cell numbers (**Fig. 1e**). Our comprehensive single-cell proteomic profiles included proteins from all subcellular localizations^22^, with a notable identification of over 200 proteins on plasma membranes, emphasizing robustness of the method (**Fig. 1f**).

### Exploring the carrier proteome effect in label-free SCP

In LFQ-based SCP, each cell is analyzed independently by LC-MS/MS with nDIA, free from the signal interferences characteristic of multiplexed SCP. However, the proteomic profiles identified from single cells can be significantly influenced by the strategy employed for the spectra-to-database matching with a peptide search engine. Two strategies are currently employed to enhance identification numbers using two popular DIA database search tools, Spectronaut^23^ and DIA-NN^24^. In Spectronaut, this is the directDIA+ or spectral library-free based approach, which includes searching alongside matched samples of higher quantities (such as 1-ng digests or those from 20 cells), whereas in DIA-NN it is the match-between-run (MBR)^25^ feature, similarly paired with higher quantity samples. While these search strategies differ between tools, both can elevate identification numbers in single cell samples by incorporating data from matched higher quantity samples. These methods are widely applied in SCP, yet their precise effects remain to be fully understood. We performed a systematic evaluation to elucidate the effect of using such a ‘carrier proteome’ for SCP by benchmarking between Spectronaut and DIA-NN, alongside their respective search strategies. When compared to searches with only single-cell samples, inclusion of carrier proteomes resulted in a substantial increase in the number of identifications (**Fig. 2a, 2b, Supplementary Figure 3a, 3b**). Notably, both DIA-NN and Spectronaut yielded nearly identical results when carrier proteomes were included in the search. Further investigation into the carrier proteome effect in Spectronaut revealed that as the number of single-cell files in a search increased, the number of identifications also rose incrementally. When carrier proteomes were included, there was a marked enhancement in identifications, increasing from around 4000 to approximately 5000 (**Fig. 2c, Supplementary Figure 3c**). Analysis of the abundance of proteins identified through different search strategies indicated that searches incorporating more cells could discern proteins of lower abundance, with the carrier proteome enabling the identification of the least abundant proteins (**Fig. 2d**). To compare the efficacy of these search strategies, we examined two separate runs from single-cell samples (**Fig. 2e**) and from 20-cell samples (**Fig. 2f**), contrasting their outcomes in DIA-NN and Spectronaut. The results showed a significant overlap in identifications, suggesting consistency across different cells and search tools. However, the protein quantification correlation between the two cells was marginally better in Spectronaut with R=0.91 compared to DIA-NN with R=0.84 (**Fig. 2g, 2h**). Conversely, the correlation between Spectronaut and DIA-NN for the same cell was not as high with R=0.79 (**Fig. 2j**).

**Figure 2.**
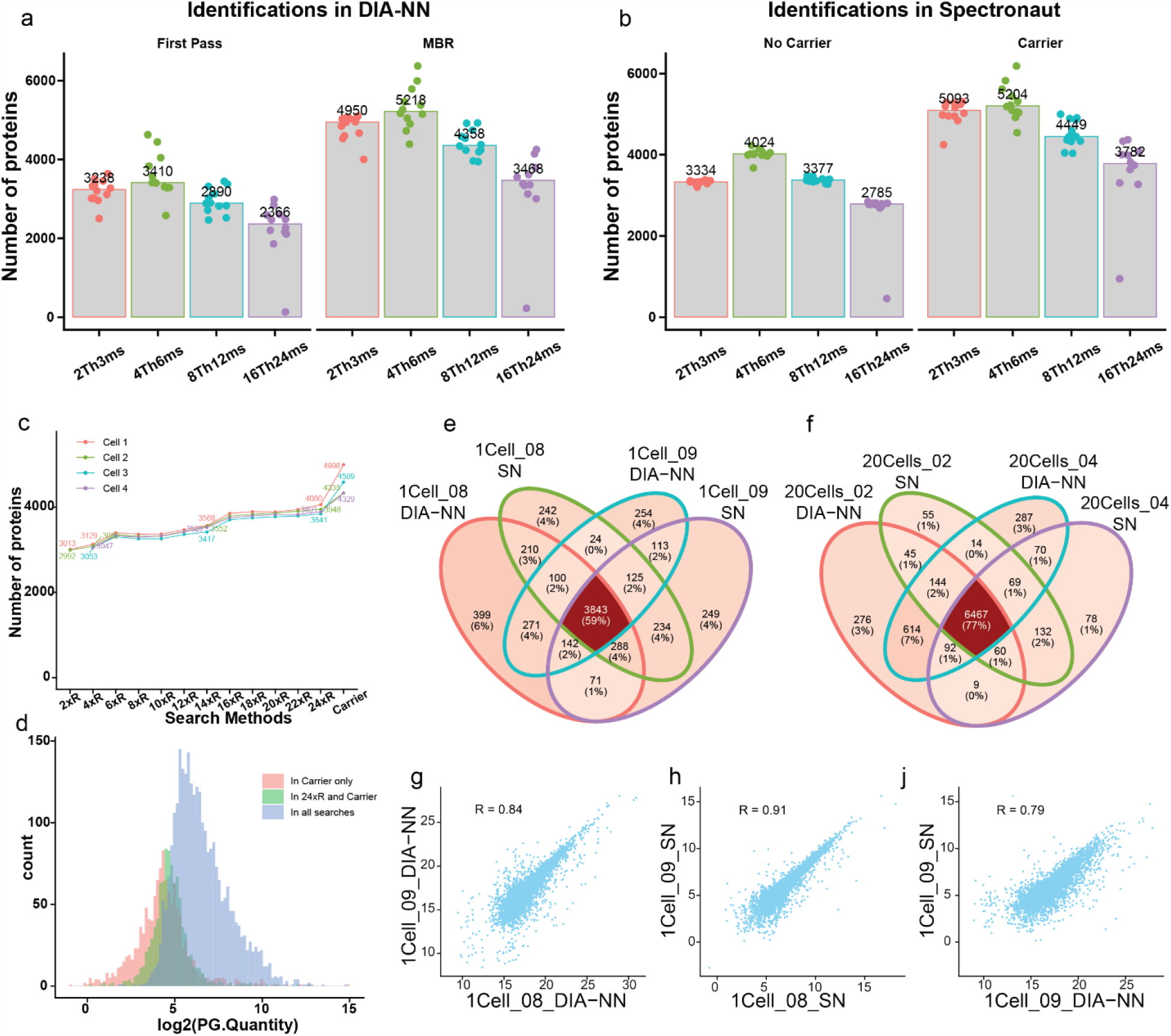
Evaluating the carrier proteome effect in single cell proteomics. (**a**) Protein identifications in single-cell samples using DIA-NN with and without match-between-run (MBR) feature, compared across various nDIA methods. (**b**) Protein identifications in single-cell samples with and without carrier proteomes in Spectronaut. (**c**) Trend analysis showing the numbers in protein identifications when single-cell data is searched with different numbers of single-cell files and with carrier proteomes in Spectronaut. (**d**) Histogram illustrating the distribution of protein abundances identified across different search strategies, highlighting the advantage of carrier proteomes in detecting low-abundance proteins. (**e**) Venn diagram comparing protein identifications in a single cell sample across two search methods in DIA-NN and Spectronaut. (**f**) Venn diagram of protein identifications comparing 20-cell samples using DIA-NN and Spectronaut, indicating the overlap and unique findings. (**g**), (**h**) Scatter plots displaying the quantification correlation of proteins identified between two single-cell runs in Spectronaut and DIA-NN, respectively. (**j**) Scatter plot presenting the quantification correlation between Spectronaut and DIA-NN for the same single-cell sample, with correlation coefficients indicating the degree of agreement. In (**e-f**), 08 and 09 are two different HeLa cells of similar sizes.

### Entrapment search analysis for error rate estimation

The ultra-low abundance of single cell proteomes raises questions about the confidence in identification, particularly with the increased numbers achieved using the Orbitrap Astral mass spectrometer. To address these concerns, we employed an entrapment database search strategy in both Spectronaut and DIA-NN to empirically estimate the error rate of identifications^26^. We initiated this by conducting an entrapment experiment with a 9x larger shuffled human mimic database, analyzing single-cell samples in combination with 20-cell samples serving as a carrier proteome with default FDR parameter settings of 0.01 at both precursor and protein level. The resulting empirical false discovery rate (FDR) at the protein level, corrected for the mimic database size, was estimated at approximately 1% (∼0.7%) (**Fig. 3a**), and less than 0.1% at the modified peptide level (**Fig. 3b**) using Spectronaut. These false identifications typically had higher protein group q-values (PG.Qvalues), i.e. low scores after negative log-transformation of the q-values (**Fig. 3c**) and displayed a random distribution in their abundances (**Fig. 3d**). In contrast, the corresponding DIA-NN analysis displayed a much higher empirical FDR when applying a PG.Q.Value threshold of 0.01 (**Fig. 3e**). However, the PG.Q.Values of these incorrect identifications were distinctly separated from the true target identifications, both in First Pass searches and MBR searches (**Fig. 3f**). Thus, applying a more stringent q-value cutoff of 0.001 could effectively eliminate false identifications with minimal impact on the overall number of true target identifications (**Fig. 3g, Supplementary Figure 4**). In summary, both Spectronaut and DIA-NN are capable of providing reliable results for single-cell samples, but it is crucial to set the q-value cutoffs to ensure data accuracy.

**Figure 3.**
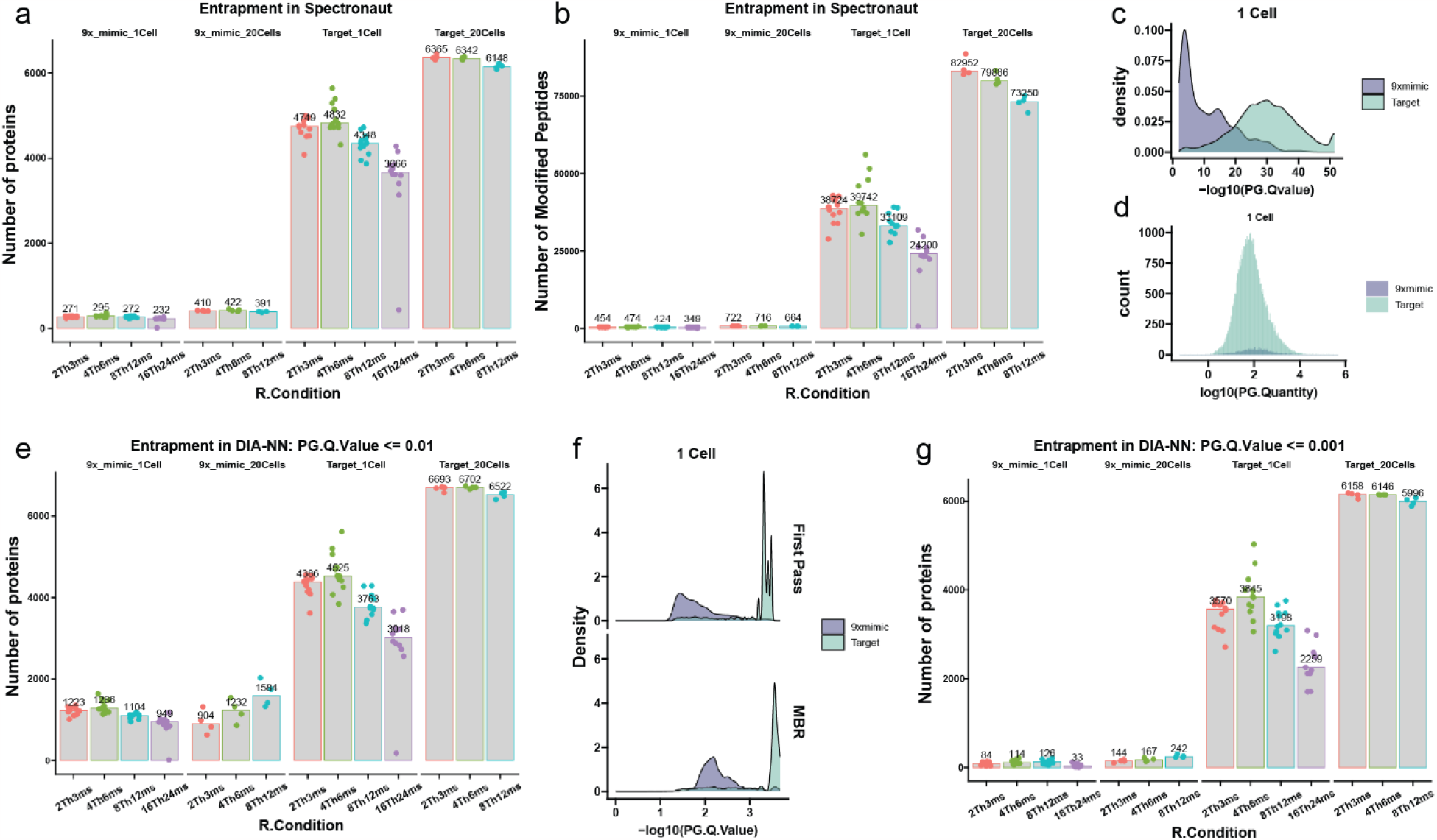
Error rate estimation in single cell proteomics through entrapment analysis. (**a**) Number of proteins identified in SCP using a 9x mimic database with Spectronaut, highlighting the empirical FDR at protein level against different nDIA methods. (**b**) Number of modified peptides identified in SCP with the 9x mimic database using Spectronaut, showcasing the empirical FDR at the modified peptide level. (**c**) Density plot for the distribution of PG.Qvalues of false and target identifications in single-cell samples using Spectronaut. (**d**) Histogram of the abundance (-log10(PG.Quantity)) of identified proteins in single-cell samples, illustrating the random distribution of false identification abundances. (**e**) Protein identification count in SCP using a 9x mimic database with DIA-NN, showing a higher empirical FDR when applying a PG.Q.Value cutoff of <= 0.01. (**f**) Density plots representing the distribution of PG.Qvalues of false identifications in single-cell samples from DIA-NN, comparing first pass and MBR searches. (**g**) Total protein identifications in single-cell and 20-cell samples using a stringent PG.Q.Value cutoff (0.001) in DIA-NN.

### In-depth PTM analysis without enrichment in SCP

The exceptional depth achieved at the peptide precursor level in our SCP datasets not only enhances protein identification and quantification but could also unveil a wealth of information pertaining to PTMs. Due to the sub-stoichiometric nature of PTMs, global analysis of any PTM by LC-MS/MS usually requires specific enrichment of the PTM-bearing peptides prior to MS analysis. However, with the high peptide coverage achieved in SCP, we speculated that identification of PTMs without specific enrichment could be possible. To test this hypothesis, we delved into the prevalence of two key PTMs of high cellular importance, phosphorylation and glycosylation, within the single-cell proteome samples, bypassing any specific enrichment processes. Focusing on the enzymes catalyzing the transfer of phosphate groups from ATP to target proteins, we quantified 168 protein kinases within single cells, encompassing all principal kinase families (**Fig. 4a**). Notably, kinases such as CDK1 from the CMGC group and MAPK1 from the STE group exhibited the highest abundances, whereas tyrosine kinases were less abundant. Subsequently, we conducted a database search for serine, threonine and tyrosine phosphorylation as a variable modification in single-cell samples, utilizing 20-cell samples as a carrier proteome. Although we did not do any specific cell perturbation or sample treatment to preserve the phosphorylation sites, the search approach led to the confident identification of a median of 120 phospho-Ser, 28 phospho-Thr, and 13 phospho-Tyr sites in single cells, with high site localization probabilities (**Fig. 4b**). A sequence logo analysis highlighted prevalent phosphorylation motifs like the SP motif corresponding to substrates for abundant kinases like CDKs, aligning closely with our kinome data (**Fig. 4c**). Importantly, an extracted ion chromatogram (XIC) screening of DIA-MS/MS spectra for the selective immonium ions for phospho-Tyr^27^ at m/z 216.042 showed intensive signals across the entire LC elution profile (**Fig. 4d**). We also investigated protein glycosylation patterns and detected multiple glycosyltransferases across all glycosylation pathways (**Fig. 4e**). Employing a strategy akin to the phospho-Tyr immonium ion screening, we performed oxonium ion screening^28, 29^ for common glycans, including the monosaccharides HexNAc (**Fig. 4f**) and NeuAc (**Fig. 4g**), and the disaccharide Hex-HexNAc (**Fig. 4h**). All glycans appeared abundantly and were present in most MS2 spectra, even when using nDIA. The screening of both immonium and oxonium ions indicated a pervasive presence of PTMs, such as protein phosphorylation and glycosylation, in single cells. Nevertheless, the precise identification of the modified peptides remains challenging due to current limitations in database search algorithms.

**Figure 4.**
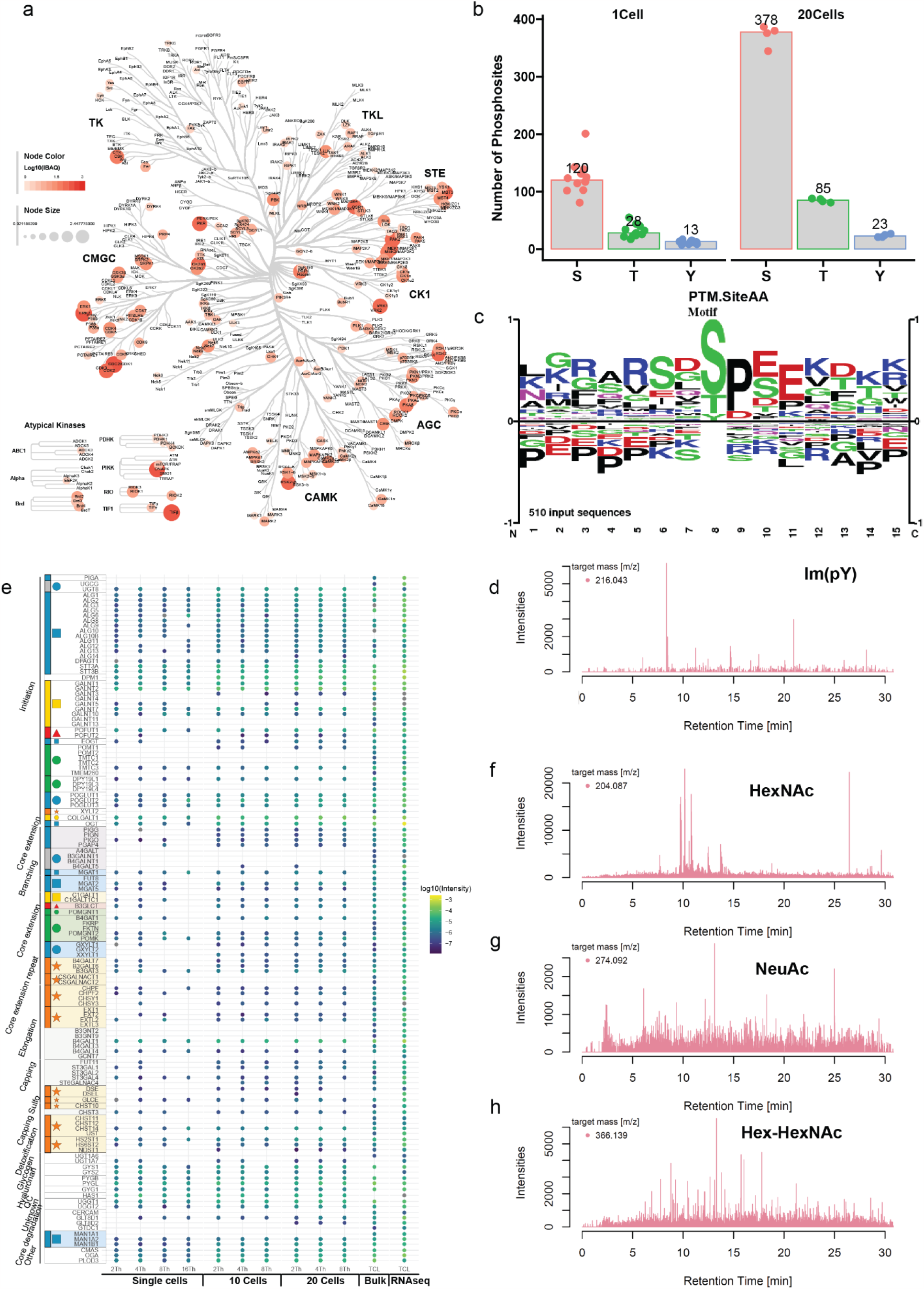
Post translational modification profiling in single cell proteomics. (**a**) Kinome tree representation showing the quantification of 168 kinases across major kinase families in single cells, with node color and size indicating iBAQ abundances. (**b**) Bar graph depicting the number of phospho-sites identified for serine (S), threonine (T), and tyrosine (Y) residues in single-cell and 20-cell samples. (**c**) Sequence logo analysis illustrating the most common phosphorylation motifs identified in single-cell samples. (**d**) Extracted ion chromatogram (XIC) of immonium ion for phosphor-Tyr (m/z 216.0426) across the LC elution profile. (**d**) Visualization of the glycogenes identified in single cells. (**f**) XIC of oxonium ion for HexNAc (m/z 204.087) indicating glycan presence in the MS2 spectra. (**g**) XIC of oxonium ion for NeuAc (m/z 274.092) reflecting glycan abundance in the MS2 spectra. (**h**) XIC of oxonium ion for the disaccharide Hex-HexNAc (m/z 366.139) reflecting glycan abundance in the MS2 spectra.

### Biological insights from single-cell spheroid analysis post 5-Fluorouracil exposure

Recognizing the throughput limitation in mass spectrometry as a significant bottleneck for single-cell proteomics, our methodology has innovated a solution that doubles the processing capacity from 40 samples per day (SPD) to an efficient 80 SPD on the Evosep One LC system. This optimized method not only enhances throughput but also maintains a high level of sensitivity, identifying over 2000 proteins in individual HeLa cells (**Supplementary Figure 5**). To demonstrate the applicability of our SCP method with 80SPD, we next intended to capture the transformative effects of the chemotherapeutic drug 5-fluorouracil (5-FU)^30^ on colorectal cancer cells grown as a spheroid. The spheroids subjected to 5-FU demonstrate increased disintegration over time, after treatment with a recently developed disaggregation buffer for 20 minutes, whereas control spheroids retained their compact structure, indicative of the impact of 5-FU on cell cohesion (**Fig. 5a, 5b**). After single cell analysis with 80SPD methods, we identified >2500 proteins in total. Conducted based on the proteome data, a PCA analysis effectively distinguished between treated and untreated single cells, which underscores the precision of the protein quantification (**Fig. 5c**). The activation pathway of 5-FU reveals an upregulation of NME1, a known 5-FU sensitizer, and a downregulation of TYMP, crucial for converting 5-FU into its DNA-incorporating active form; these changes are crucial indicators of drug efficacy^31^ (**Supplementary Figure 6**). The hierarchical clustering of gene ontology terms underscores the involvement of biological processes such as cyclase cytoplasmic activity and ribose purine synthesis, which are directly affected by mechanism of 5-FU action on rRNA synthesis (**Fig. 5d**). The representation of these pathways aligns with the known impact of 5-FU on the nucleotide synthesis pathways. An upset plot of the altered gene ontology terms further dissects the effects of 5-FU, detailing which processes are most affected by the treatment and their regulatory trends (**Fig. 5e**). The alteration in proteins associated with these gene ontology terms highlights specific proteins like ADCY, which contributes to pyrophosphate formation, and keratin, integral for spheroid structural integrity. These findings suggest a targeted disruption by 5-FU on spheroid stability and purine metabolism, a reflection of the ability of the drug to interfere with key cellular functions (**Fig. 5f**).

**Figure 5.**
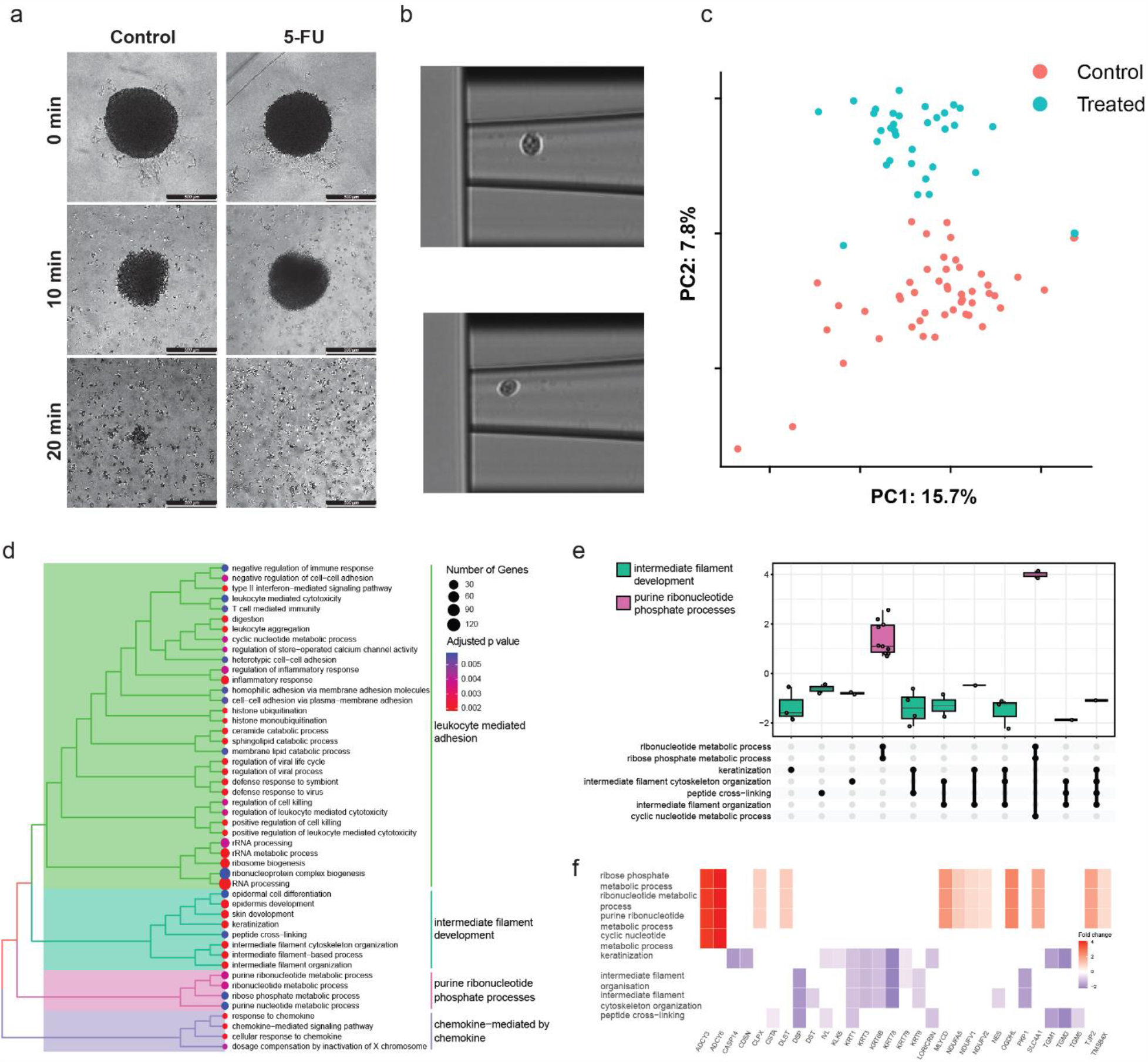
Analysis of 5-Fluorouracil impact on spheroid cells using SCP. (**a**) Comparative images illustrating the morphological changes in spheroids untreated and treated with 5-FU, showing increased disintegration and cell detachment in treated spheroids over time. (**b**) Examples of two cells being dissociation and isolated in cellenONE. (**c**) PCA analysis of the single cells untreated and treated with 5-FU. (**d**) Hierarchical clustering of Gene Ontology terms related to cellular processes affected by 5-FU treatment, highlighting the involvement of cyclase cytoplasmic activity and ribose purine synthesis in its action mechanism. (**e**) UpSet plot of the top altered Gene Ontology terms post-5-FU treatment, detailing the upregulated and downregulated biological processes. (**f**) Heatmap of protein alterations across identified Gene Ontology terms.

## Discussion

Our study marks a significant leap in the field of single-cell proteomics, achieving a remarkable enhancement in sensitivity, with identifications soaring from approximately 2000 proteins to a staggering 5000 proteins. This breakthrough underscores the evolution of the powerful proteomics technologies empowering researchers to delve deeper into the molecular intricacies of individual cells. The combination of lossless single-cell sample preparation using the cellenONE with the ultra-high sensitivity provided by the Evosep One Whisper flow gradients and nDIA on the Orbitrap Astral is a powerful setup enabling SCP with high coverage and robustness. The rigorous evaluation of software tools and the implementation of FDR strategies were pivotal in bolstering the confidence of our protein identifications. Through meticulous analysis, we discerned the subtleties and strengths of various computational approaches, ensuring the accuracy of our findings and reinforcing the validity of our data. Notably, we demonstrated the feasibility of PTM analysis without the prerequisite of specific enrichment protocols. This advancement paves the way for a more streamlined and efficient exploration of PTM landscapes, revealing the multifaceted regulatory mechanisms at play within the cell. The introduction of a spheroid-specific dissociation buffer has also proven influential, enhancing the dissociation efficiency of spheroids, and bolstering the robustness of our proteomic analysis. This innovation stands as a testament to the relentless pursuit of methodological refinement in SCP.

Despite these advancements, we recognize that the throughput in mass spectrometry is a major bottleneck, currently capped at 80 samples per day. We propose that this limitation could be mitigated through the integration of nDIA with multiplexed approaches, including TMT and multiplexed DIA. This combinatorial strategy holds the potential to exponentially increase throughput and analytical depth. Meanwhile, exciting progress has been made in non-MS techniques for identifying and potentially sequencing individual proteins^32, 33^. These methods draw inspiration from nucleic acid sequencing technologies and include single-molecule peptide sequencing through Edman degradation or amino peptidases in flow cells, as well as the use of nanopore sequencing adapted for proteins.

Our workflow is poised to become a cornerstone in future SCP studies, illuminating the technological advancements and applications in complex dynamics of cellular function and disease. Looking to the future, we foresee the integration of single-cell genomic, proteomic and other omics measurements as a transformative approach. We are on the verge of an era where multi-omics experiments will become the norm, and their synergistic application promises to provide a more comprehensive understanding of cellular states, particularly in disease contexts.

## Methods

### Cell lines

Different human cell lines (HeLa) were cultured in DMEM (Gibco, Invitrogen), supplemented with 10% fetal bovine serum, 100U/ml penicillin (Invitrogen), 100 μg/ml streptomycin (Invitrogen), at 37 °C, in a humidified incubator with 5% CO2. Cells were harvested at ∼80% confluence by washing three times with PBS (Gibco, Life technologies). Cells were then resuspended in degassed PBS at 200 cells/μL for isolation within the CellenONE®.

### Spheroid formation

Prior to spheroid formation HCT116 cells were seeded to P15 plates and grown to a confluence of 70-90%. Subsequently, cells were washed with Phosphate-buffered saline (PBS) (Gibco, Life Technologies) and detached from the plate with Trypsin. Following this the cells were counted using a Corning® Cell Counter (Sigma-Aldrich). For the multicellular spheroid generation 7.000 cells were seeded on ultra-low attachment 96-well plates (Corning CoStar, Merck). The spheroids were cultured for 72h at 37 °C, in a humidified incubator with 5% CO2. Cell medium was refreshed after 48 h, by aspirating half the old medium (making sure not to alter the spheroid) and adding the same amount of fresh medium. Subsequently the spheroids were treated for 24h with 2 uM 5-Fluorouracil (previously identified as the IC50 for this cell lines). After 24 hours the spheroids were transfered to an eppendorf tube using a p1000 and washed thrice with ice cold PBS. The spheroids were disaggregated using the new dissociation buffer (Cellenion SASU) by incubating for 15 minutes shaking at RT followed by 10 minutes shaking at 4 °C. Additionally the eppendorf tube was gently shaken to induce mechanistical disintegration as well. Following this 10 μl of cell solution was diluted in 490 μl of cold PBS prior to sample preparation in the CellenONE®.

### Sample preparation in cellenONE and proteoCHIP EVO 96

Sample lysis and digestion is performed within the proteoCHIP EVO 96 inside the CellenONE. This process begins with the manual deposition of 2 μL of oil into each well of the chip, which is then positioned on the target plate. The entire system is cooled to 8°C to ensure the oil solidifies effectively. Following this, a master mix consisting of 0.2% DDM (D4641-500MG, Sigma Aldrich, Germany), 100mM TEAB, 20 ng/μL trypsin, and 10 ng/μL lys-C in a volume of 300 nl is dispensed into each well.

The cell isolation process utilizes the precision of the CellenONE module to sort individual cells, which are selected based on a diameter range of 22 - 30 μm and a maximum elongation factor of 1.6, into the wells. The proteoCHIP is then subjected to a controlled incubation phase at 50°C with 85% relative humidity for 1.5 hours within the instrument’s environment. After incubation, the system’s temperature is reduced to 20°C to stabilize the conditions post-reaction.

Upon completion of the incubation, the proteoCHIP EVO 96 is taken out of the CellenONE and processed further. This involves the addition of 4 μL of 0.1% formic acid (FA) to each well, followed by a chilling period at 4°C to re-freeze the oil. Parallelly, the Evotips are prepared in line with the vendor’s guidelines, which include a series of rinsing, conditioning, and equilibrating steps with specified solvents to prepare them for sample uptake. Afterward, another 15 μL of 0.1% FA is introduced to each tip, and the proteoCHIP is promptly inverted onto the Evotips and centrifuged at 800g for 20 seconds at 4°C.

Once the proteoCHIP EVO 96 is removed, the Evotips undergo a customized procedure that deviates slightly from the standard vendor’s protocol. The samples are first loaded onto the Evotips by centrifugation at 800 g for 60 seconds, ensuring that the peptides are fully captured by the tip matrix. After loading, the Evotips are meticulously washed with 20 μL of Solvent A, and the centrifugation step is repeated for another 60 seconds at 800 g to remove any non-specifically bound substances, thereby increasing the purity of the captured peptides. The final step involves adding 100 μL of Solvent A to the Evotips and spinning them for a brief 10 seconds at 800 g. The samples are then ready for LC-MS/MS analysis.

### LC-MS/MS

The LC-MS/MS analysis was conducted using an Orbitrap Astral Mass Spectrometer coupled with an Evosep One chromatography system from Evosep Biosystems. The sample runs were set up for either a Whisper 40SPD (31-minute gradient) or Whisper 80SPD (15-minute gradient) method. We utilized Aurora Elite TS analytical columns from IonOpticks with an EASY-Spray™ ion source. The Orbitrap Astral operated with a resolution setting of 240,000 for full MS scans across a mass-to-charge range of 380 to 980 m/z. The automatic gain control (AGC) for full MS was adjusted to 500%. MS/MS scans employed various narrow-window data-independent acquisition (nDIA) methods with isolation windows and ion injection times tailored to the specificity of the analysis—these included settings like 2Th3ms, 4Th6ms, 8Th12ms, and 16Th24ms. The MS/MS scanning spanned the same m/z range of 380 to 980.

Fragmentation of the isolated ions was carried out using Higher-energy Collisional Dissociation (HCD) set at a normalized collision energy (NCE) of 27%. For the 40SPD method, single-cell samples were distributed among the different nDIA methods: 12 for 2Th3ms, 12 for 4Th6ms, 24 for 8Th12ms, and 12 for 16Th24ms. Samples composed of 10, 20, and 40 cells were analyzed in sets of four using the 2Th3ms, 4Th6ms, and 8Th12ms methods respectively.

### Data analysis

For the analysis using Spectronaut v18 (Biognosys), raw files underwent a library-free directDIA+ approach, employing the human reference database from the Uniprot 2022 release, which contains 20,588 sequences, alongside an additional 246 common contaminant sequences. Notably, cysteine carbamidomethylation was not included as a modification, while variable modifications were set for methionine oxidation and protein N-terminal acetylation. The precursor filtering was based on Q-value, and cross-run normalization was not applied. For the phosphorylation search, phosphorylation on serine, threonine and tyrosine was set as a variable modification.

In the case of DIA-NN searches, the raw data files were first converted to .mzML format. The analyses were then conducted in a library-free mode with some modifications: the maximum number of variable modifications was limited to two, and like with Spectronaut, cysteine carbamidomethylation was not set as a modification. It was configured to employ highly heuristic protein grouping, and the match-between-runs (MBR) feature was activated.

For the entrapment analysis, entrapment peptides were processed using an updated version of a tool known as “mimic” from the percolator repository. This tool shuffles the target database ninefold and appends the shuffled proteins flagged as “mimic” and “Random_XXXX”, preserving the original amino acid composition from the fasta file. This strategy ensures a rigorous assessment of the false discovery rates in our proteomic analysis.

