## Supplementary Figures for "High-throughput and scalable single cell proteomics identifies over 5000 proteins per cell"

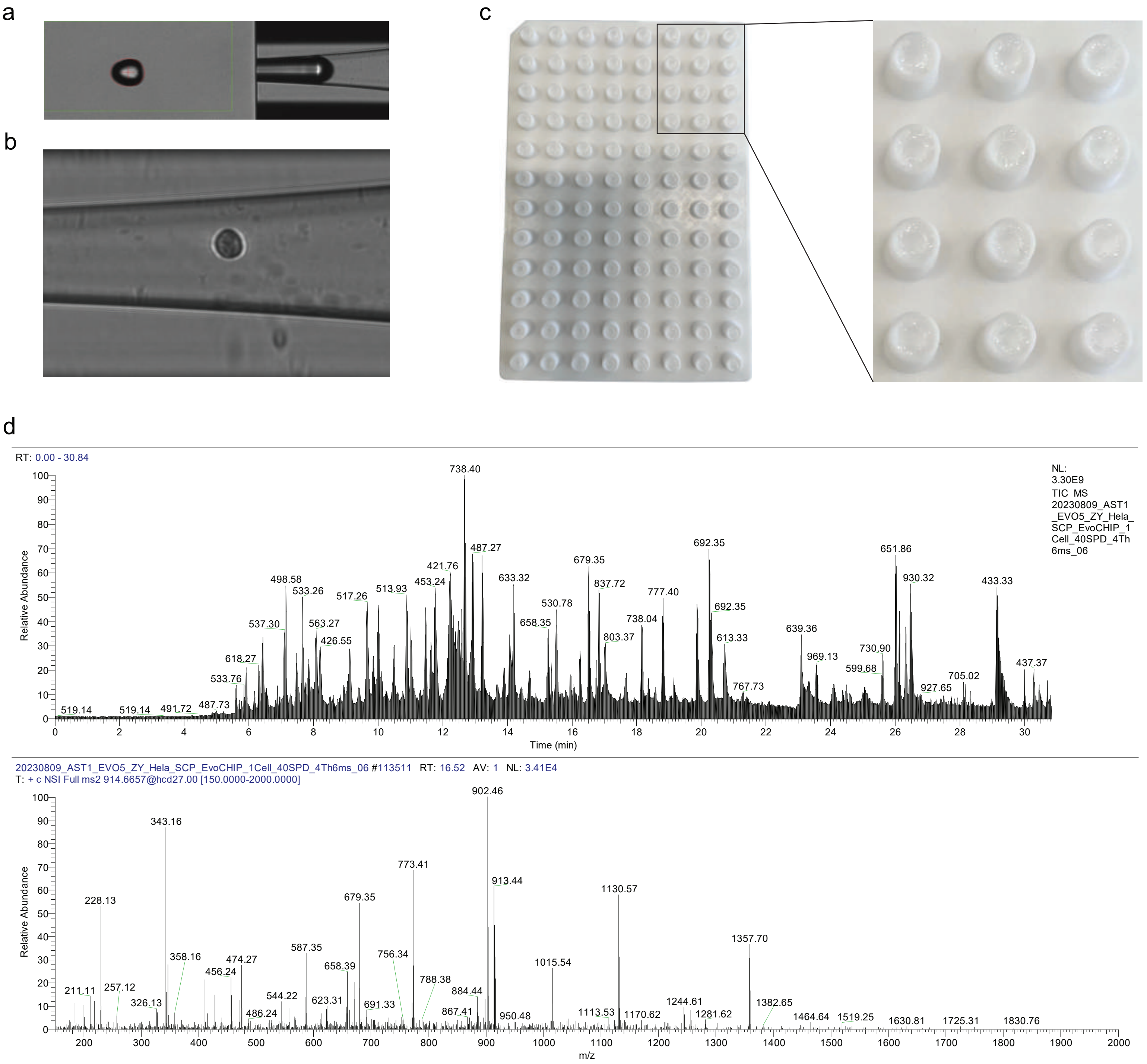


**Supplementary Figure 1 | Examples of each step in the SCP workflow.**

a) Example of a droplet in cellenONE. This is used in buffer dispensing.

b) Example of an isolated HeLa cell in cellenONE.

c) Picture of a proteoCHIP EVO 96. This is the picture taken after the sample transfer from the chip to Evotips.

d) Example of a total ion current (TIC) and an MS/MS spectrum in a single cell sample.

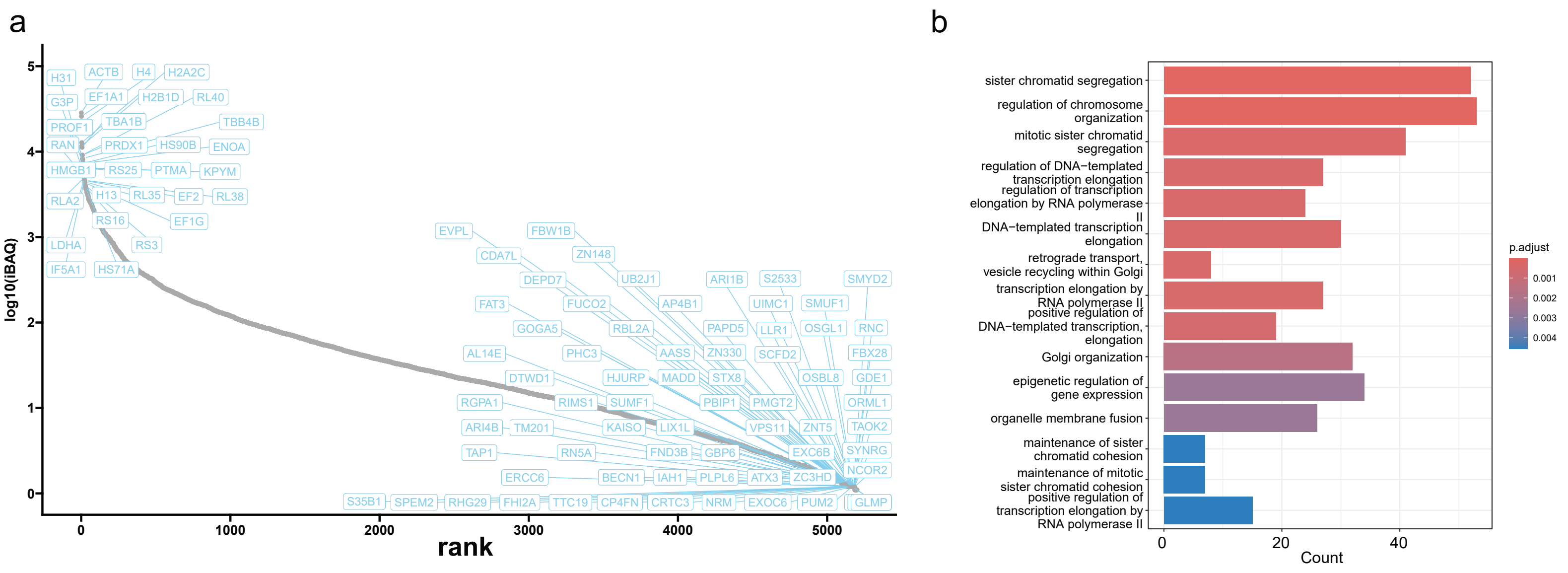


**Supplementary Figure 2 | iBAQ distribution of the single cell proteome.**

a) Distribution of the iBAQ values a single cell sample, with labels indicating some of the highest and lowest abundant proteins.

b) Geene ontology enrichment analysis specifically focusing on the 1500 proteins with the lowest abundance.

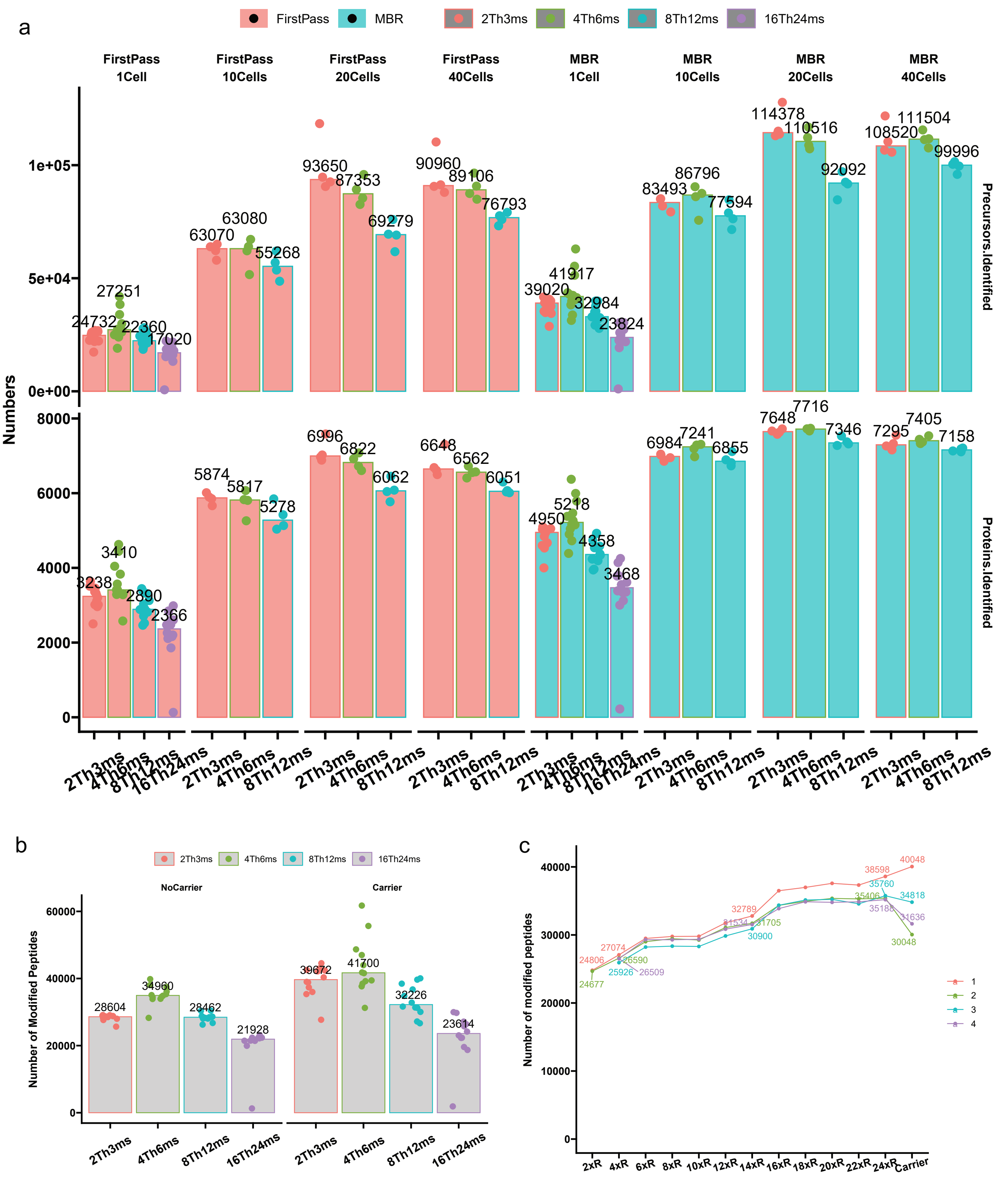


**Supplementary Figure 3 |Carrier proteome effect in label-free single cell proteomics.**

a) Number of proteins and precursors identified in the HeLa samples in DIA-NN.

b) Number of modified peptides identified in the single cell HeLa samples in Spectronaut.

c) Trend analysis showing the numbers in peptide identifications when single-cell data is searched with different number of single-cell files and with carrier proteomes in Spectronaut.

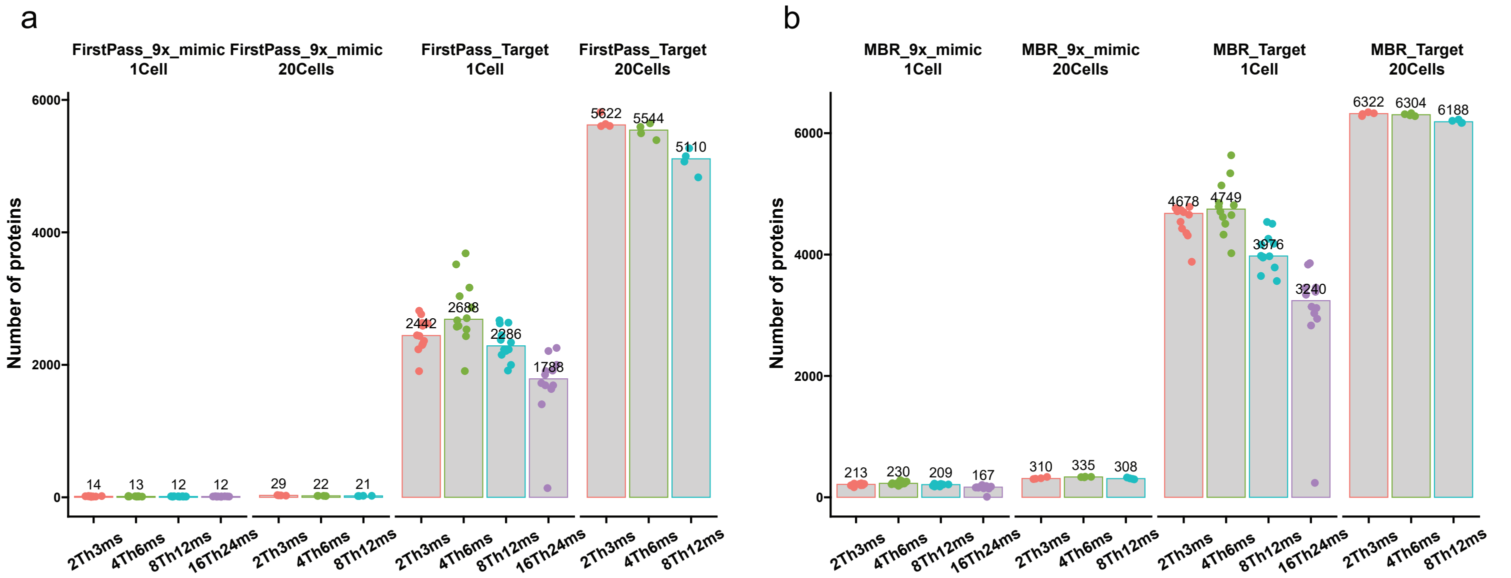


**Supplementary Figure 4 | Total identifications in single-cell and 20-cell samples using a stringent Global.PG.Q.Value cutoff (0.0005) in DIA-NN with first search approach (a) and MBR approach (b).**

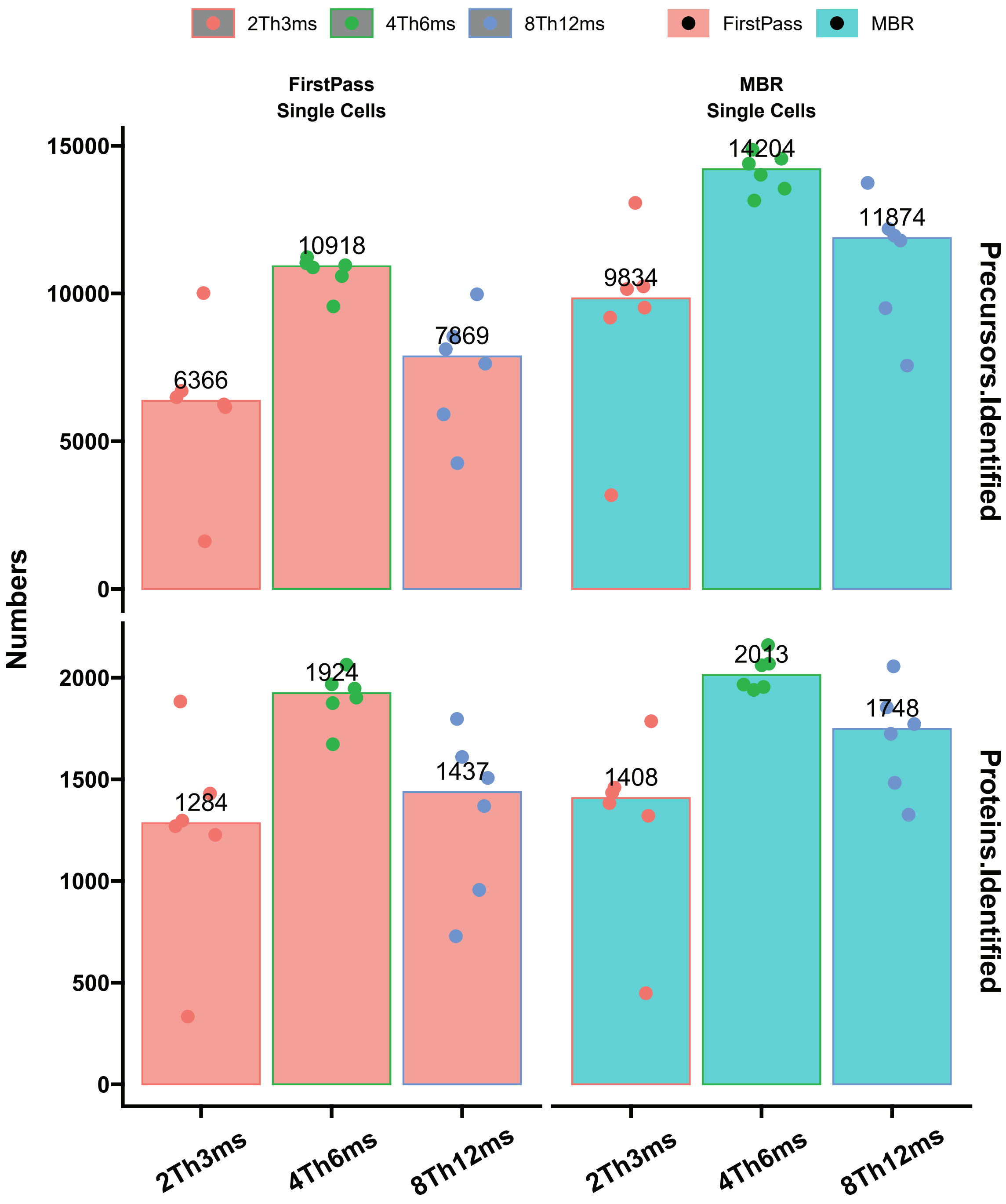


**Supplementary Figure 5 | Number of proteins and precursors identified in the HeLa samples run with 80 SPD method.**

6 single cell samples were analyzed in each method. The search was done in DIA-NN.

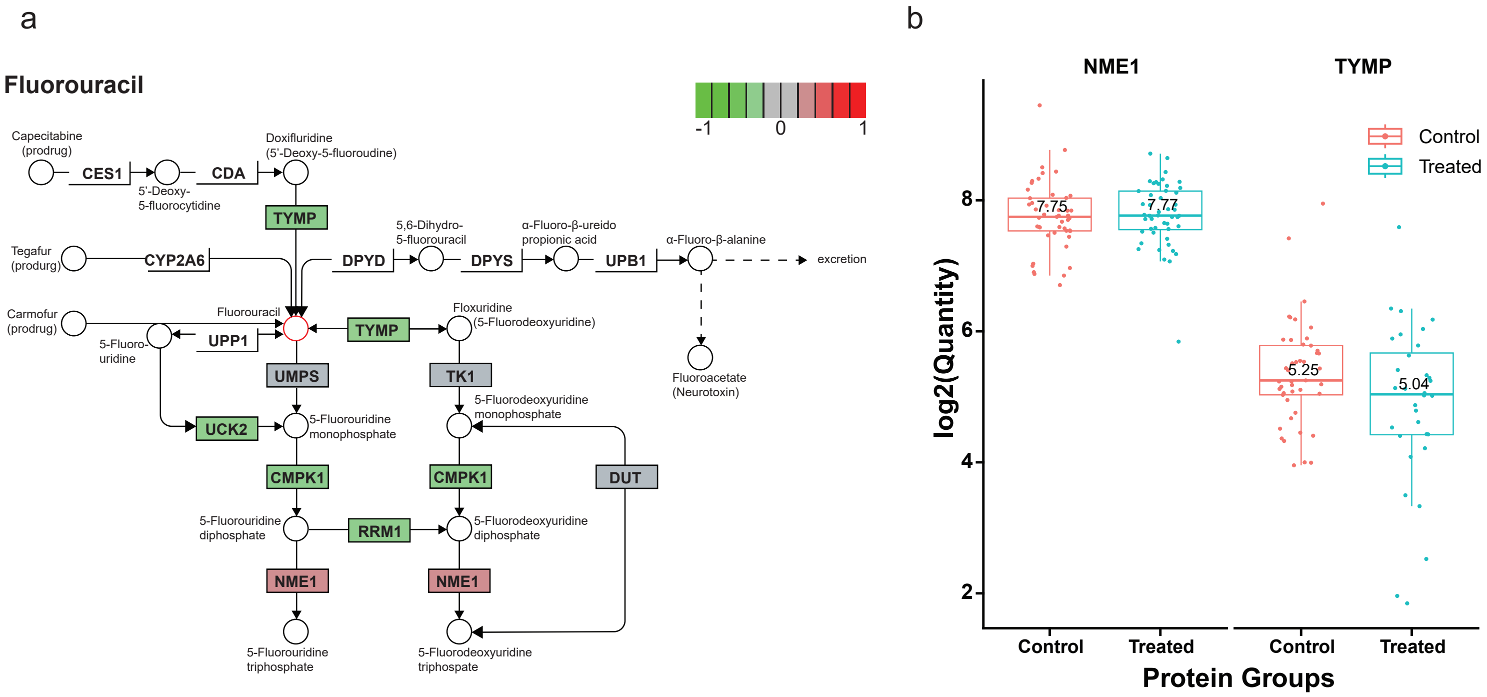


**Supplementary Figure 6 | iBAQ distribution of the single cell proteome.**

a) Metabolic pathway diagram of 5-FU indicating the regulation of NME1 and TYMP, proteins integral to the metabolism and activation of 5-FU.

b) Abundance distribution of TYMP and NME1 in the single cells.
